# Multi-Scale Structural Analysis of Proteins by Deep Semantic Segmentation

**DOI:** 10.1101/474627

**Authors:** Raphael R. Eguchi, Po-Ssu Huang

## Abstract

Recent advancements in computational methods have facilitated large-scale sampling of protein structures, leading to breakthroughs in protein structural prediction and enabling *de novo* protein design. Establishing methods to identify candidate structures that can lead to native folds or designable structures remains a challenge, since few existing metrics capture high-level structural features such as architectures, folds, and conformity to conserved structural motifs. Convolutional Neural Networks (CNNs) have been successfully used in semantic segmentation — a subfield of image classification in which a class label is predicted for every pixel. Here, we apply semantic segmentation to protein structures as a novel strategy for fold identification and structural quality assessment. We represent protein structures as 2D α-carbon distance matrices (“contact maps”), and train a CNN that assigns each residue in a multi-domain protein to one of 38 architecture classes designated by the CATH database. Our model performs exceptionally well, achieving a per-residue accuracy of 90.8% on the test set (95.0% average accuracy over all classes; 87.8% average within-structure accuracy). The unique aspect of our classifier is that it encodes sequence agnostic residue environments from the PDB and can assess structural quality as quantitative probabilities. We demonstrate that individual class probabilities can be used as a metric that indicates the degree to which a randomly generated structure assumes a specific fold, as well as a metric that highlights non-conformative regions of a protein belonging to a known class. These capabilities yield a powerful tool for guiding structural sampling for both structural prediction and design.

**Significance:** Recent computational advances have allowed researchers to predict the structure of many proteins from their amino acid sequences, as well as designing new sequences that fold into predefined structures. However, these tasks are often challenging because they require selection of a small subset of promising structural models from a large pool of stochastically generated ones. Here, we describe a novel approach to protein model selection that uses 2D image classification techniques to evaluate 3D protein models. Our method can be used to select structures based on the fold that they adopt, and can also be used to identify regions of low structural quality. These capabilities yield a powerful tool for both protein design and structure prediction.

## Introduction

While the 3D structures of proteins arise as a consequence of their amino acid sequences, computational design methods operate by creating sequences that minimize the energy of a pre-defined protein backbone. This approach has provided solutions to challenging design problems ranging from enzyme catalysis to viral inhibition(1–8). Computational design begins with defining a set of constraints that constitute the design problem, followed by either acquiring a natural backbone (from the Protein Data Bank) or building one from scratch (*de novo* design). Once a backbone is defined, amino acid sequences are designed onto it(9).

Obtaining a suitable backbone is the most challenging step in computational protein design. Highly evolved native backbones tolerate only minimal deviations from their original sequences, leading to a restricted set of solutions for a given design problem. *De novo* design, which uses physical and chemical principles to build proteins from scratch, offers at least three advantages over native-based methods(10). First, because the role of each residue in a *de novo* structure is precisely defined, the range of tolerated modifications is well understood, allowing for more controlled customization. Second, *de novo* proteins often have greater thermodynamic stability, which facilitates the introduction of function(11, 12). Lastly, by not being restricted to existing structures, *de novo* backbones offer solutions otherwise unattainable from native scaffolds, are likely more adaptable to a wider range tasks(10).

Despite these advantages, *de novo* design is challenging because it requires the construction of a protein with neither a known structure nor sequence. Since the vastness of the protein torsion space (φ, ψ, χ) prohibits an exhaustive search, current *de novo* design protocols generate backbones by combining small, continuous segments of φ-ψ torsions (“fragments”) collected from the Protein Data Bank (PDB). While this allows for a significant reduction in search space, an enormous amount of sampling is still required to find the lengths, types (e.g. helix, beta-sheet) and order of secondary structure elements (here on, “topologies”) that result in viable structures. The process of identifying successful models and topologies currently relies on a combination of scoring functions, hydrogen-bonding patterns, and other discernible regularities to screen for models that satisfy the desired criteria(13–19). However, such heuristics are often subjective and chosen *ad hoc*, and there are currently no unbiased, generalizable methods that can perform automated structure selection based on the overall organization of a protein.

Meanwhile in the field of computer vision, convolutional neural networks (CNNs) have revolutionized pattern-recognition tasks ranging from facial recognition(20) to object detection(21). In applications to proteins, several groups have used 1D and 3D CNNs to process protein sequence and structure data, performing tasks such as domain prediction(22) and mutation stability evaluation(23). Nonetheless, several limitations have prevented the use of these models in protein design. In the case of 1D CNNs, low-dimensionality often results in loss of important features needed to describe realistic structures. In the case of 3D CNNs, model sizes (i.e. the number of weights in a network) scale quickly with input size, resulting in significant hardware requirements for efficient processing of full-length protein structures. Most 3D representations also lack rotational and translational invariance, thus requiring large quantities of data to train deep models. Surprisingly, few reported studies have used 2D CNNs to perform protein structure analysis, despite the fact that 2D CNNs are the most well-studied and widely implemented class of neural network.

In this study, we demonstrate how CNNs intended for 2D image processing can be used as powerful 3D structure analysis tools to guide computational protein design. A number of recently reported 2D CNN architectures have enabled advanced forms of image classification, namely instance(24) and semantic(25) segmentation, which predict class labels for individual pixels as opposed to entire images — segmentation provides information about where exactly an object is in an image, rather than simply indicating whether or not an object is present. We speculated that this capability is pertinent to protein structure analysis and hypothesized that image segmentation networks could be adapted to create a model that quantifies structural features at various scales within a protein (e.g. per-residue, per-domain).

We trained a 2D CNN that classifies each residue in a protein into one of 38 architecture classes designated by the CATH(26) protein database **(Figure 1a)**. We represent a protein using the pairwise distance matrix (here on, “contact map”) between all α-carbons in its structure(27, 28) **(Figure 1b)**. This representation is rotationally and translationally invariant, and provides an image-like rendering of 3D protein structures in 2D. In the same way that image segmentation predicts a class for the basic unit of an image -- a pixel, our model predicts classes for the basic unit of a protein -- a residue.

**Figure 1:**
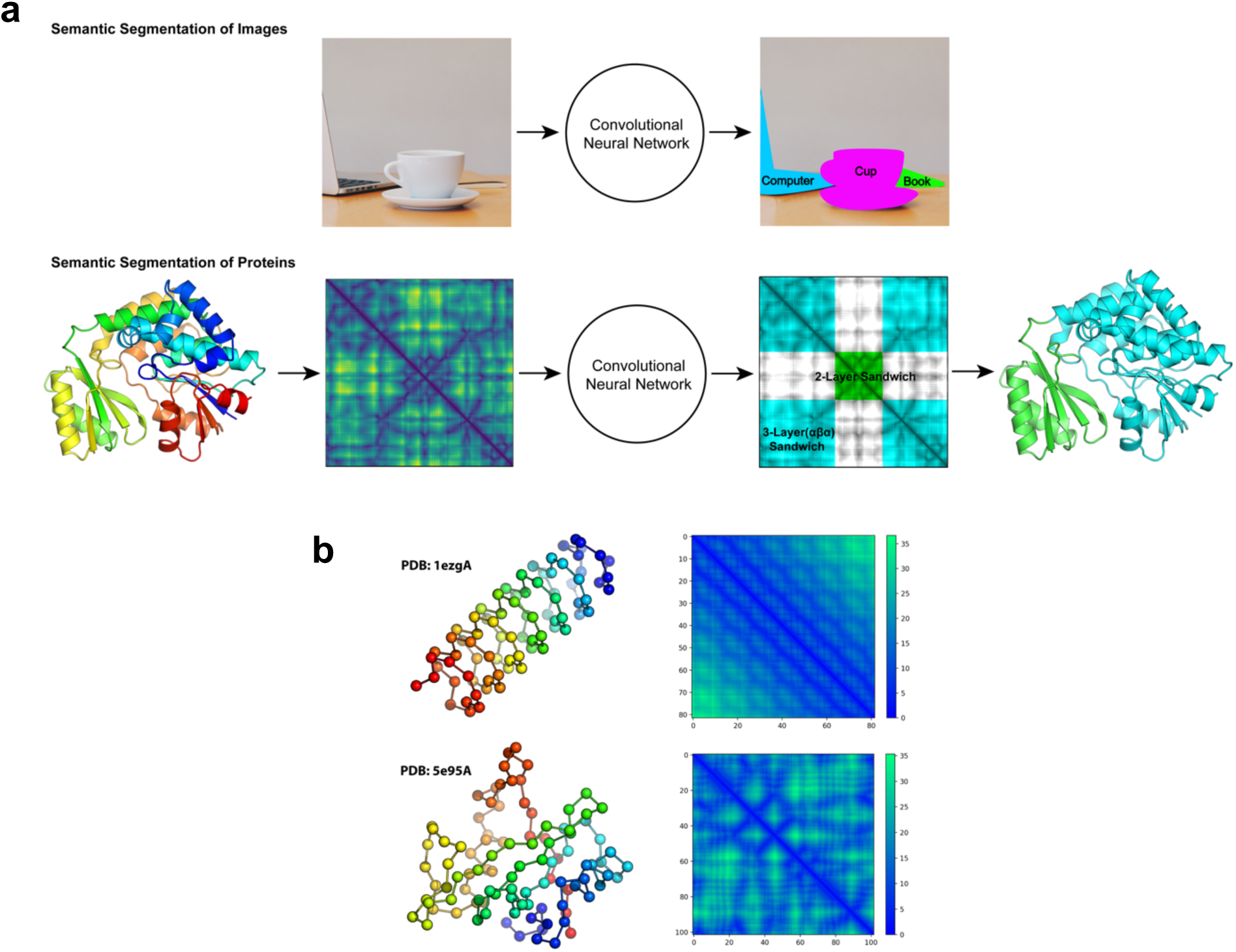
Semantic Segmentation of Protein Structures using Contact Maps. **a**, A comparison of semantic segmentation for objects in images, and domains in proteins. The two-domain protein shown (PDB ID: *1mlaA*) contains a 3-layer-αβα-sandwich (cyan), and a 2-layer sandwich (green). Multiple distal regions in a contact map can correspond to a single domain in a protein. The example segmentation and contact maps are real inputs and outputs obtained from our model. **b**, Two examples of single-domain proteins colored as chainbows (left), with α-carbons shown as spheres. Red indicates the beginning, and blue indicates the end of the chain. The corresponding contact maps are shown to the right of each protein with units in Å. Axes correspond to amino acid index.

While the most obvious function of our network is in protein classification, its true significance lies in its utility as a *de novo* design tool. On the single-residue level, Shannon entropies(29) of the CNN-predicted probability distributions can be used as indicators of local structure quality, allowing for identification of low-confidence regions that require further refinement or reconstruction. On the full-protein level, per-residue class probabilities predicted by our classifier can be averaged across an entire protein to provide a quantitative measure of the degree to which a structure assumes a fold. This function is useful for quickly searching large design trajectories for proteins that adopt specific architectures.

## Results

Our motivation for using a per-residue classifier as a protein evaluation tool is based on the assumption that the environment of each residue is rich in structural information(30), which can be aggregated to describe higher level features such as domains boundaries, architectures, and structure quality. Importantly, the goal of our study is not to create a protein classifier, but rather to show how a classifier can be used to quantify residue-level environments in the form of probability distributions, which can then be averaged over regions of a protein to obtain information at the desired size scales. Our classifier thus differs from past protein classification methods(26, 31) in that it performs per-residue domain-architecture classifications based only on distance information and is completely sequence agnostic. In the *Supplementary Information*, we describe how our classifier can be extended to perform conventional protein classification.

### Performance as a Per-Residue Classifier

Class predictions for a row (or column) in a contact map are effectively a prediction in 3D space, because each row can be mapped to the location of a residue in the original structure. Using this fact, we trained a CNN to perform semantic segmentation on proteins, which entails predicting one of 38 architectural classes for each residue. In the same way that image segmentation models assess the context of individual pixels in an image, our CNN evaluates local features in the contact map, which encode structural features centered around a residue in 3D space.

A diagram of our CNN is shown in **Figure 2a**, with additional technical details provided in the *Methods* section. The network converged within 100 epochs of training **(Figure. 2b)** and achieves average test accuracies of 90.8% per residue, 95.0% per architecture class, and 87.8% per structure. Because the proportion of the dataset comprised by each class is highly variable, one concern is that the model may have simply learned to predict frequently occurring classes. **Figure 3a** suggests otherwise, as there is no correlation between the frequency of a class in the training set and class accuracy in the test set. The majority of class accuracies are above 90% even for the rarest classes **(Figure 3a and 3b)**.

**Figure 2:**
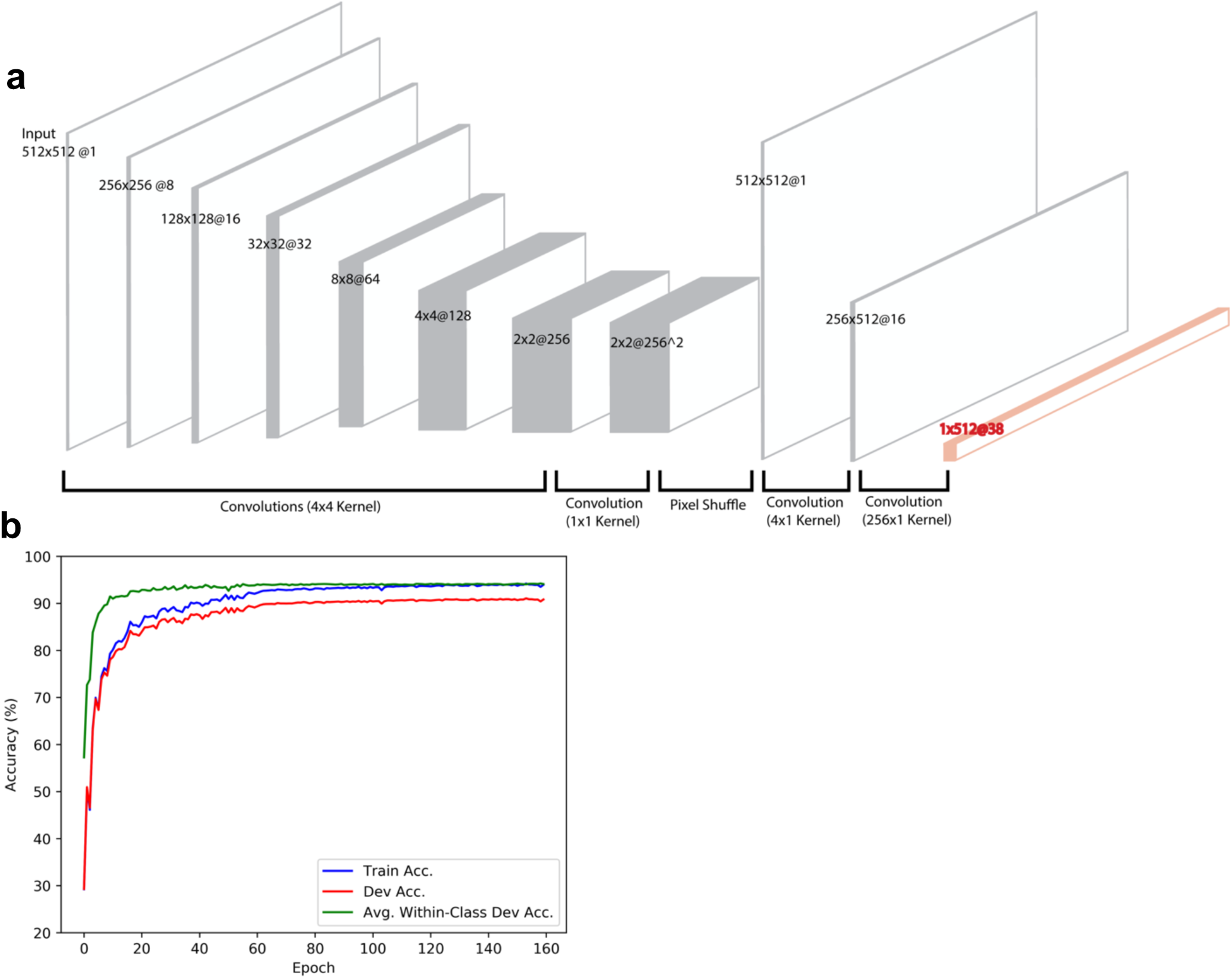
Model Architecture and Training. **a**, The model is comprised of six down-sampling convolutional layers (encoding), followed by a “pixel shuffle” step for up-sampling to a 512×512 map (decoding). Rectangular convolutions are used to reshape the feature map into a 1×512@38 which corresponds to a probability distribution over the 38 CATH architecture classes for each residue (orange). **b**, Accuracy profiles of the training and development sets during training. The model converges within 100 epochs, achieving a residue-wise training accuracy above 90%. The performance generalizes to the test set, where the model achieves a per residue accuracy of 90.8%, a per-class accuracy of 95.0%, and a per-structure accuracy of 87.8%.

**Figure 3:**
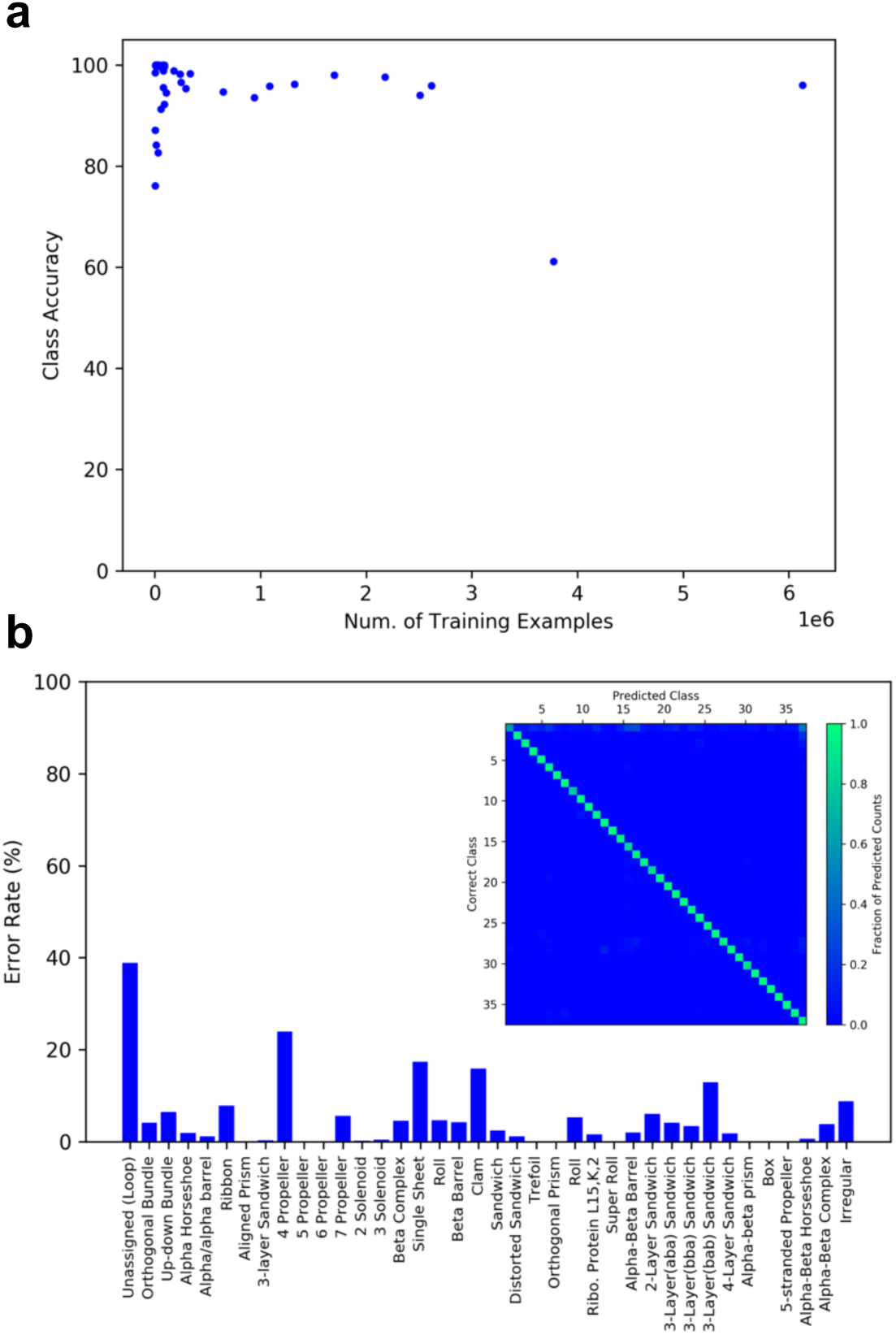
Analysis of Classifier Performance. **a**, A plot of the number of residues in the training set belonging to a specific class, versus the class accuracy in the test set. Each point in the plot represents a single class. No obvious correlation between the number of examples in the training set and class accuracy on the test set is observed. The majority of class accuracies are above 90% even for the rarest classes. **b**, Error rates for each architecture class. The confusion matrix is shown as an insert with architectures indexed in the same order as on the horizontal axis of the bar graph. The vertical axis of the confusion matrix indicates the correct class. The horizontal axis indicates the class predicted by the model. Each column is normalized to the total number of predicted counts for each class.

Example outputs (PDB IDs: *3gqyA, 3sahB, 4i1eA*) are shown in **Figure 4**. Overall the model is able to closely reproduce per-residue architecture classifications provided by the CATH database. Inspection of outputs suggests that our model is able to accurately classify individual residues, even if they constitute fragments of a domain that are isolated in sequence space. This is illustrated in the case of the central αβ-barrel in *3gqyA* **(Figure 4, 3gqyA classifications, cyan)**, which is comprised of multiple fragments from distal regions in primary sequence (**Figure 4, 3gqyA, chainbow**). This observation suggests that the network is able to correlate delocalized features in the 2D contact map corresponding to the same domain in the 3D structure. Additionally, our network is able to recognize differences in secondary structure organization. For example, both the orthogonal-bundle in *3sahB* **(Figure 4, *3sahB,* green)** and the α-horseshoe in *4i1eA* **(Figure 4, *4i1eA,* green)** are purely alpha-helical in content, and both are comprised of residues in contiguous sequence. Although these architectures differ only in their secondary structure organization, the model is able to successfully differentiate between the two.

**Figure 4:**
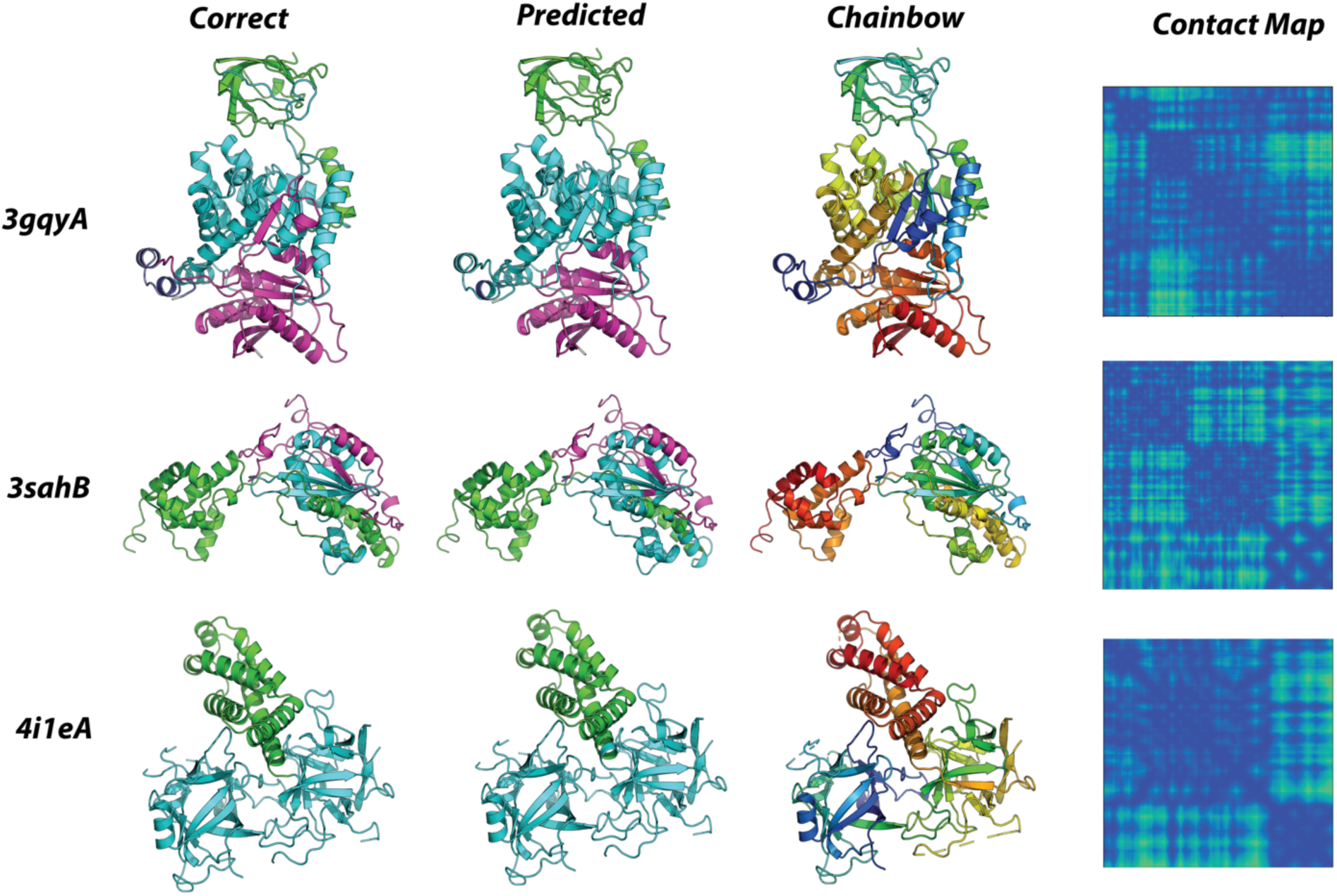
Classifications and Corresponding Contact Maps. Segmented examples from the test set. From the left, each column depicts: the correctly segmented structure, the model prediction, protein chain colored in residue order (rainbow order), and the input contact map. PDB ID’s for each protein are shown on the left. *3gqyA*: 86.9% within-structure accuracy: beta-barrel (green), alpha-beta barrel (cyan), loop (dark blue), 3-layer-αβα(magenta). *3sahB*: 89.0% within-structure accuracy: orthogonal-bundle (green), 2-layer sandwich (cyan), loop (magenta). *4i1eA*: 99.2% within-structure accuracy: alpha-horseshoe (green), trefoil (cyan).

A characteristic of our model that distinguishes it from most conventional structure classification algorithms(26, 31) is that it is sequence-agnostic. This is particularly important for *de novo* proteins, which have sequences that lack homology with naturally occurring families. The *de novo* designed TIM barrel structure, sTIM-11 (PDB ID: *5bvl*), is one such example(13). sTIM-11 belongs unequivocally to the αβ-barrel architecture class, however it remains unclassified in both CATH(26) and SCOPe(31), the two largest protein classification database, likely due to its lack of sequence homology to native TIM barrels. Our model is able to correctly classify this structure.

### Generalizability of Learned Structural Features

Although our model is able to achieve high classification accuracies, accuracy metrics alone provide little information about the generalizability of the features it has learned. Given that neural networks are highly flexible, it is possible that our classifier is fitting to features such as protein length, which may correlate with architecture class but provide little structural information. To control for such possibilities, we tested whether the features encoded by our classifier could generalize to other structure evaluation tasks (*viz*. transfer-learning(32)). Specifically, we trained a “small” 6-layer CNN and assessed its ability to predict the Global Distance Test Total Score(33) (here on, GDT) of CASP submissions(34). The input to the small network is the 1×512@38 layer of the classifier network, which encodes the probability distributions over the 38 architecture classes at each residue (**Figure 2a, orange**) — this is effectively a reduced representation of a protein, encoded by our classifier **(Figure 5a)**.

**Figure 5:**
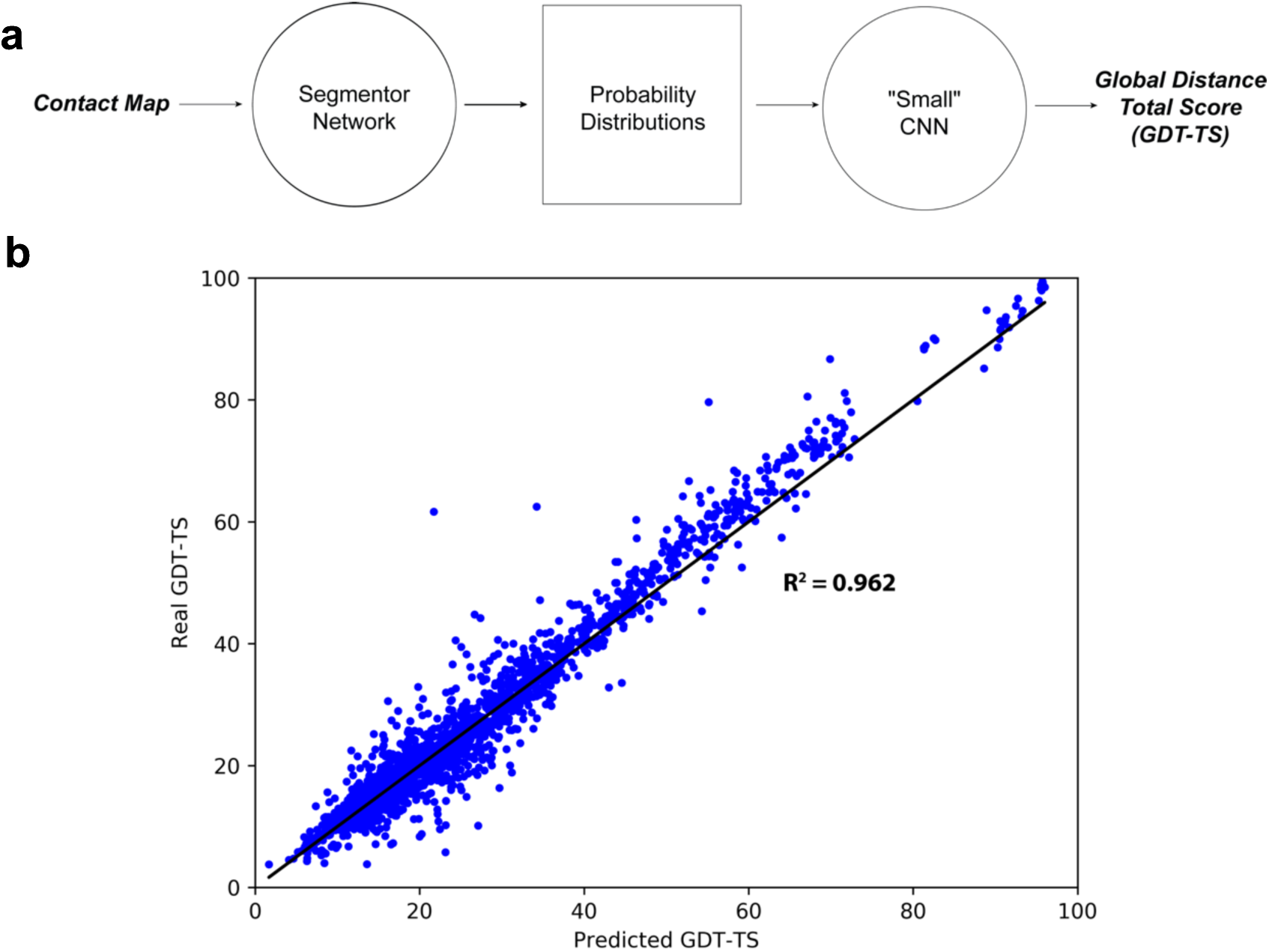
GDT-TS Predictions for 1,960 submissions for 98 CASP Target Domains. **a**, A schematic of the GDT-TS prediction scheme. **b**, A plot of real GDT-TS vs the GDT-TS predicted by the small model. The identity function is shown as a black line. Each point corresponds to a single decoy from the test set, which consists of 20 randomly chosen submissions for each of the 98 targets (1,960 total decoys). The small model was able to accurately predict the GDT-TS of the CASP submissions, yielding an R-squared value of 0.962 between the real and predicted values. The 98 selected targets were exclusively in the free-modeling categories of CASP10~12, and were not in the training set of the classifier model.

The small network was trained on 41,029 CASP submissions for 98 target domains exclusively in the free-modeling categories of CASP 10-12, and that were not contained in the training set of the classifier network. 20 randomly chosen submissions from each of the 98 targets were reserved for use in the test set. The small CNN model was able to accurately predict the GDT of the CASP submissions, yielding an R-squared value of 0.962 between the real and predicted GDT values **(Figure 5b)**. Plots of the real versus predicted GDT values for each target are provided in the **Supplementary Figure S6**.

While the predictive power of the small model is limited to the 98 targets, these results suggest that the representation learned by the classifier is rich in structural information at sub-Å resolution. The accurate predictions despite that many of the CASP targets did not fall into any CATH architecture class, suggesting that the learned structural features are useful for describing any protein, not just those belonging to the predefined classes.

### Selecting Valid Design Topologies

A significant challenge in *de novo* design is that it is necessary to find a viable set of lengths, types, and orders of secondary structure elements that can adopt a target fold. “12-residue α-helix, 3-residue loop, 5-residue β-strand” is one such example, and is referred to as a “topology.” In shorthand we denote this topology as “H12-E5”, where “H” and “E” denote helix and sheet, while numbers denote element lengths.

To test whether our CNN can be used to perform topology selection, we generated a series of design trajectories targeting the TIM-barrel fold (αβ-barrel architecture). Our previous study discovered that a four-fold repeat of E5-H13-E5-H11, with 3-residue loops interspersed between each secondary structure element, constitutes a viable TIM-barrel topology(13). In order to test whether our model could distinguish this validated topology from a pool of random ones, we generated backbones for every permutation of the secondary structure elements in the four-fold E5-H13-E5-H11 repeat while maintaining the 3-residue loops between each of the helix or sheet elements; this resulted in 12 unique topologies. We used *RosettaRemodel*(35), which was used in the original study, to generate 7000 backbone designs for each of the topologies, producing a total of 84,000 structures.

For each candidate structure we used our network to predict the average probability of the αβ-barrel architecture over all residues (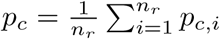 where p_c,i_ denotes the probability of class *c* at residue *i*). We observed that the predicted probabilities correlated with the degree to which a structure adopts a TIM-barrel fold (**Figure 6a**). The distribution of probabilities for the generated structures exhibits a bimodal distribution with only ~2% of structures receiving a probability prediction greater than 50% **(Figure 6b)**. Collecting the top 2000 designs corresponding to the high probability peak, we observed that the majority of the recovered structures (1301 models, 65%) belonged to 5E-13H-5B-11H topology, suggesting that our model was able to identify the experimentally validated topology as being the most promising. The proportions of the topologies found in the top 2000 structures is shown in **Figure 6c** along with the highest probability structures from each. Designs from highly represented topologies tended to have fewer structural irregularities relative to under-represented ones. The top four topologies shared the common feature of having two helical elements separated by a single E5 element, likely due to the fact that the having the second E5 element at either the beginning or end of the repeating unit results in near-equivalent structures that are circular permutations of one another. It has been experimentally confirmed that circularly permuted TIM-barrel variants fold correctly(13). These results suggest that our model can be used in combination with established protein backbone generation protocols, such as *RosettaRemodel*, to automatically select viable protein topologies.

**Figure 6:**
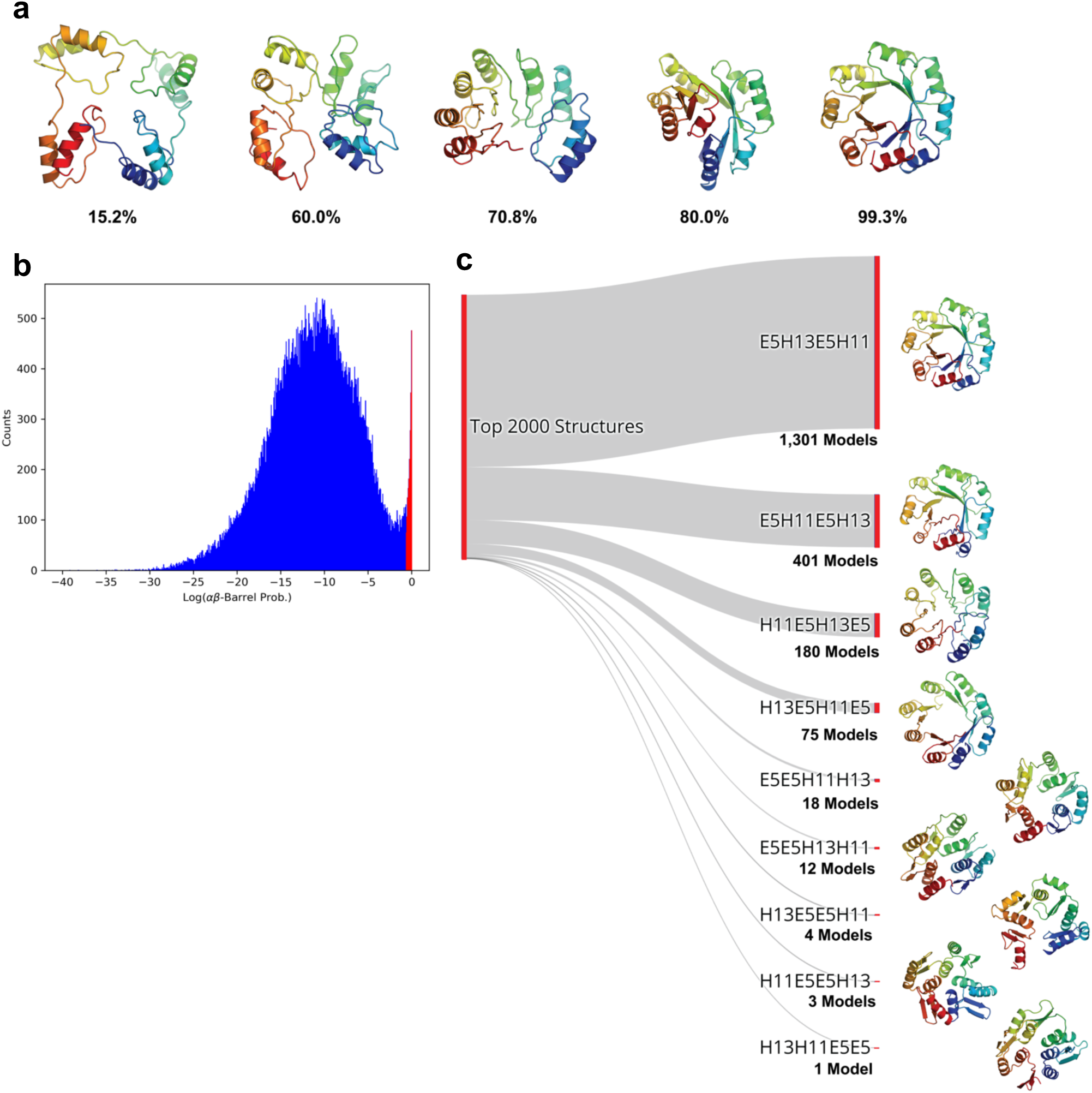
Identification of Viable TIM Barrel Topologies. **a**, Models generated in the *de novo* design trajectory and their predicted αβ-barrel probabilities. **b**, The distribution of αβ-barrel architecture probabilities for 84,000 structures spanning 12 topologies. The region corresponding to the 2,000 structures described in **c** is shown in red. **c**, The proportions of the topologies found in the top 2,000 highest αβ-barrel probability structures. The experimentally validated 5E-13H-5B-11H topology constituted 1,301 models (65%) of the top scoring structures. The highest probability structure from each topology is shown to the right.

### Precision and Local Structure Evaluation

While the result above suggests that our model may be useful for design, it does not provide insight into the sensitivity of our metric. Because unviable topologies largely produce models that differ significantly from their target architecture, the task of identifying promising topologies is a coarse-grained problem. To stringently test whether our classifier can quantify the degree to which a structure adopts a particular fold, we assessed our network’s ability to differentiate between highly similar models, using predicted class probabilities and the Shannon entropies of the per-residue probability distributions. We test our classifier on structure prediction data because these trajectories generate a diverse ensemble of structures that converge towards a known structure -- a subset of these structures are highly similar and therefore provide a good test of model performance.

We generated a structure prediction trajectory for the PDB structure *2chfA* using the *Ab Initio* protocol(36) of Rosetta. *2chfA* was not included in the training set of the classifier, and has an αβα-3-layer-sandwich architecture. For each generated decoy we computed the probability of the αβα-3-layer-sandwich class averaged over all residues. With increasing RMSD from the native structure, the model predicted a decreasing probability of belonging to the αβα-3-layer-sandwich class **(Figure 7a)**. Importantly, the predicted probability of a class was indicative of how consistent the features of a decoy were with the features of a target fold, independent of its RMSD to the native structure **(Figure 7b)**. Predicted αβα-3-layer-sandwich probabilities correlated with the formation of a well-defined central 5-stranded β-sheet, suggesting that our model correctly identifies this β-core as a feature of this fold. These results suggest that the predicted class probabilities of our network can be used as continuous metrics for selecting decoys assuming a specific fold.

**Figure 7:**
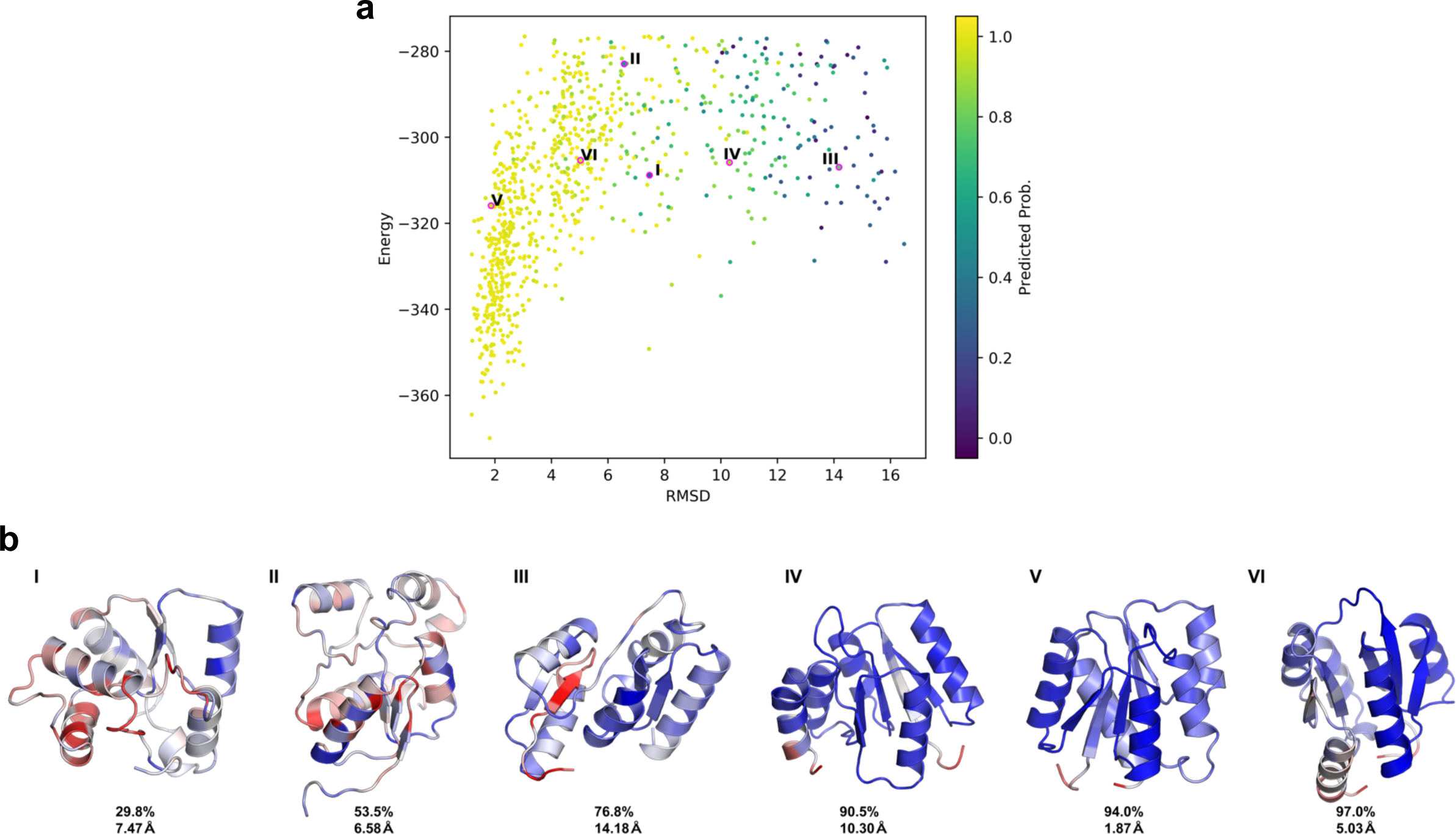
Analysis of a Structure Prediction Data. **a**, 850 randomly selected structures from a structure prediction calculation generated by Rosetta *Ab Initio*. The vertical axis indicates the energy of each structure as predicted by Rosetta. The horizontal axis indicates the RMSD of each decoy relative to the native structure. Each point is colored by probability for the αβα-3-layer-sandwich architecture, averaged over all residues in the decoy, as predicted by the classifier model. The model generally predicts a decreasing probability of belonging to the αβα-3-layer-sandwich class with increasing RMSD. Roman numerals correspond to decoys shown in **b**. **b**, Six examples from the *2chfA* structure prediction calculation. The average predicted probability of the αβα-3-layer-sandwich class, and RMSD relative to the native *2chfA* structure is shown below each decoy. The predicted probability of a class was indicative of how closely a decoy resembled the target fold, independently of native RMSD. The structures are colored by the exponential of the entropies of the predicted probability distributions at each residue. The coloring is normalized within each structure; for each decoy, red indicates the highest entropy point, and blue indicates the lowest entropy point.

The entropy *H* of the class probabilities at each residue (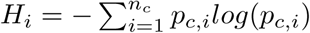), is a measure of the certainty inherent to each prediction. We observed that the high entropy regions of decoys tended to be structurally ambiguous or deviant from the target fold. This effect is shown in **Figure 7b**, where the normalized spread of entropy values within each structure is shown with a blue-to-red color spectrum. In the high probability models (**Figure 7b, IV~VI**), high entropy regions corresponded to loops, and in **Figure 7b, III** the network correctly identifies the interface of unpaired beta strands failing to form a properly assembled core as a low confidence region. In **Figure 8**, we show two other examples in which entropy appeared to be indicative of local model quality. The *de novo* structure in **Figure 8a** was generated using a αβ-2-layer-sandwich topology described by Koga et. al.(14), and the authors report that atom-pair constraints between the two C-terminal β-strands are required to obtain properly annealed designs. In an unconstrained trajectory, our model identifies the interface of the unpaired strands as being low confidence (**Figure 8a, red)**, agreeing with this statement. The structure shown in **Figure 8b** was generated in a structure prediction calculation for the gene *NHLRC3*, which has an unknown structure thought to be a 6-blade-β-propeller. In the model shown, each of the six blades has formed, however one of the blades has failed to anneal to the rest; our model identifies this sixth blade as being a low-confidence region (**Figure 8b, red)**.

**Figure 8:**
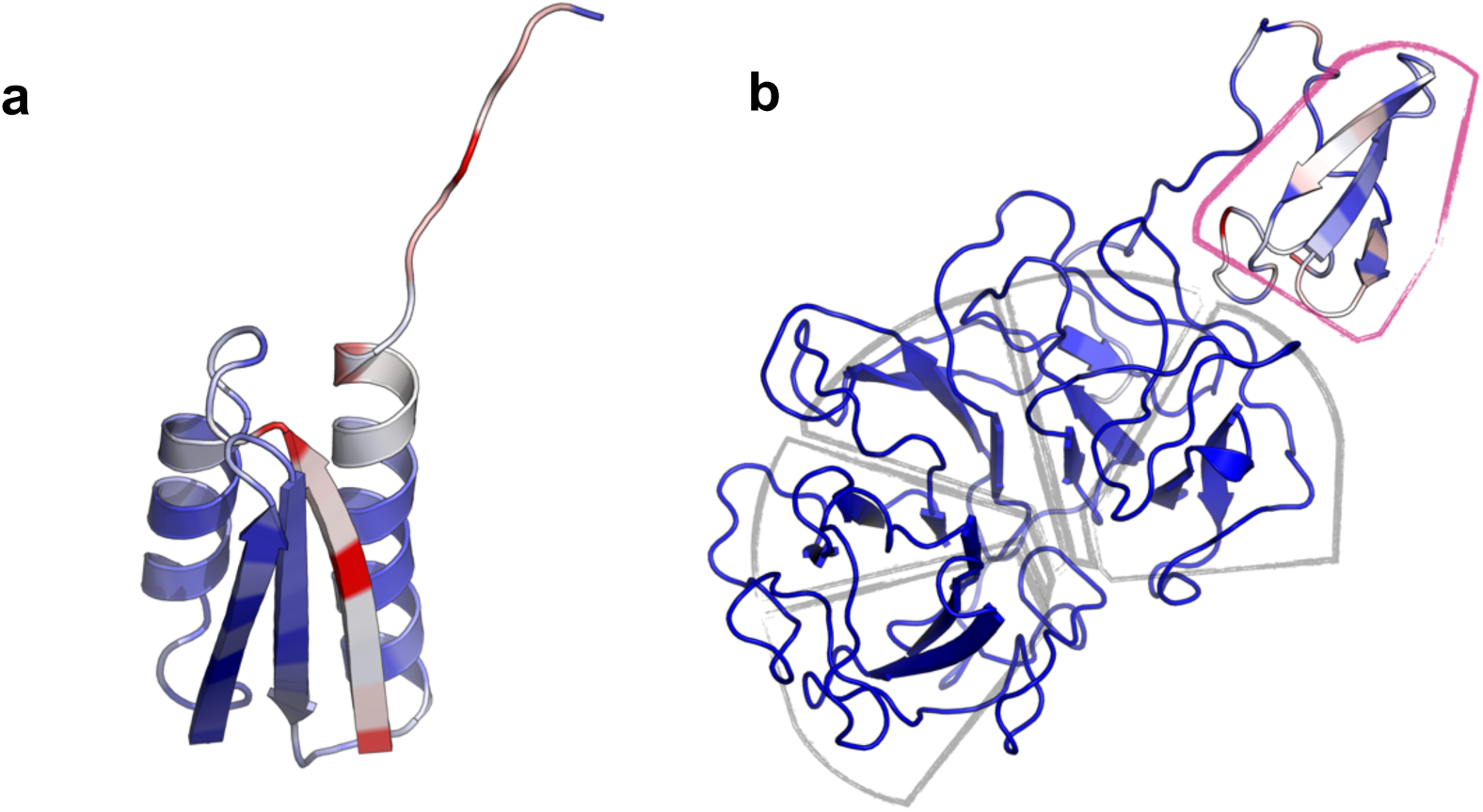
Entropy as an Indicator of Local Model Quality. Two structural models colored by the exponential of the entropies of the predicted probability distributions at each residue. The coloring is normalized within each structure; for each decoy, red indicates the highest entropy point, and blue indicates the lowest entropy point. **a**, A *de novo* structural model generated using an αβ-2-layer-sandwich topology described by Koga et. al. (2012). The interface of an the unannealed β-strands is identified as a high-entropy region. **b**, A structural model generated in a structure prediction calculation for the gene *NHLRC3*, which has an unknown structure thought to be a 6-blade-β-propeller. Five of the propeller blades have assembled (grey outline), and the un-assembled sixth blade is identified as a high-entropy region (magenta outline).

## Discussion

In recent years, new computational tools have allowed sampling of protein structural conformations at unprecedented scales, leading to breakthroughs in structure prediction(37, 38) and the creation of *de novo* designed structures -- an achievement that was considered impossible just 30 years ago(10). Nonetheless, for both prediction and design, finding a target structure requires large-scale sampling of conformational space before a subset of models with the desired features can be found. For structure prediction calculations, promising models can be selected based on criteria such as degree of convergence(39) and score function evaluations(36) because native structures fold into a minimal-energy state dictated by their sequences(40). For *de novo* design, however, the objectives of a sampling calculation are far less clear as there is no longer a sequence to guide the search process. To identify valid structures, designers have relied heavily on custom-made heuristics based on hydrogen-bonding patterns, geometric parameterizations, modified scoring functions, and other artificially defined constraints. Despite these efforts, a generalizable model selection strategy that captures high-level features such as topology, and secondary structure organization has yet to be developed.

Previously reported methods have shown that recurring structural features (motifs) can be used to accelerate conformational searches and assist in structure evaluation(30, 41, 42). We have developed a novel approach that uses a CNN to capture and quantify such features. Our CNN can be implemented in established protocols to significantly improve sampling efficiency in both structure prediction and design: first, the Shannon entropies of per-residue probabilities can be used as a measure of local model quality. Methods such as *Iterative Hybridize*(43), which iteratively recombines candidate models to obtain improved structures, could benefit greatly from using this type of method to identify low quality regions requiring rebuilding and refinement. Second, full-structure probabilities can be used to quantify the degree to which a structure assumes a target fold; this can be used to efficiently screen for models adopting a desired architecture in large sampling pools. Finding satisfactory models is comparable to finding a needle in a haystack, and is a daunting task in *de novo* design. Our novel model selection method can be used to improve the sampling efficiency by focusing searches around viable candidates in structural space. In contrast to the custom heuristics described above, our classifier is unbiased in that it is informed by 95% of all unique structural data, and generalizes to the 38 architecture classes that comprise this dataset. Our CNN is also fast, and can compute both entropies and full-structure probabilities in milliseconds, efficiently processing the millions of candidate structures generated during both prediction and design.

There are two minor limitations to our network. First, albeit by choice, inputs to our current model are limited to single chains that are 512 residues or less (95% of all unique chains in the PDB). For analysis of longer chains, structures need to be fragmented or truncated. Nonetheless, this limitation can be easily remedied by training a new model with a larger input layer. Second, our network is trained on the set of architectures defined by CATH. While CATH classifications account for the vast majority of observed protein structures, this implies that protein designers seeking to build unclassified architectures cannot currently benefit from our model.

Nonetheless, the novel way in which we apply image segmentation methods to protein structure allows us to quantify abstract structural patterns constituting the “environment” of each residue. Our results suggest that by combining per-residue information in different ways, it is possible to create any number of metrics that describe various characteristics of a protein; this approach is powerful in its generalizability and can be used to encode features of proteins that are often difficult to assess when building structural models.

## Methods

### Dataset

Training a CNN for semantic segmentation requires large amounts of densely labelled data, which are often expensive and or difficult to obtain. In the case of proteins, this corresponds to a large collection of proteins with a class assignment for each residue in every protein. Databases such as CATH(26) and SCOPe(31) provide this information in the form of residue-level domain boundary assignments and classifications. The dataset used in this study was obtained from CATH v4.2, which is comprised of 132,380 unique, fully annotated protein chains (42 architecture classes, 202,506 domains, 3.3×10^7^ residues). From this set, chains shorter than 520 residues that did not contain domains belonging to classes with fewer than 10 members were selected for use. Structures larger than 512 were center-cropped, and smaller structures were zero-padded to 512. The resulting dataset contains 126,069 chains (38 architecture classes, 181,753 domains, 2.9×10^7^ residues) spanning 95% of all unique chain data. Selection did not drastically alter the overall structure of the data **(Supplemental Figure S1a-c)**. Of the selected chains, 8,000 were reserved for each the test and development sets and the remaining 110,069 were used in training. The dataset is highly imbalanced with the largest class containing nearly 7 million residues and the smallest containing less than 3,500. To address this, the split was performed in a stratified manner, but stochastically adjusted so that all sets had at least 650 residues from each class present **(Supplemental Figure. S1d-f).** During training, examples were weighted to ensure that each class had equal influence.

### Model Architecture and Implementation

Our architecture is comprised of six convolutional layers (“encoding”), followed by a “pixel-shuffle” step(44) for upsampling to a 512×512 feature map (“decoding”). Rectangular convolutions are then used to reshape the map into a 1×512@38 tensor (**Figure 2a, orange**) that is passed to a Softmax function(32). Each convolutional layer in the encoding phase is followed by a BatchNorm(32) and a LeakyReLu(32). The 4×1 convolution is followed only by a LeakyReLu, and the final layer passes directly into the Softmax function.

The model was implemented in the PyTorch deep learning framework(45). Training was performed for a total of 160 epochs with a mini-batch(32) size of 64 using the Adam optimization algorithm. A learning rate of 0.001 was used for the first 60 epochs, 0.0001 for an additional 100 epochs. The loss function is a cross entropy loss(32) averaged across every residue in the input chain. Importantly, the loss was weighted so that training examples from under-represented classes had increased influence on training. Weights were chosen so that each class, in total, had equal influence. Dropout regularization(32) was used throughout the convolutional layers in the encoding phase with a zeroing-probability of 0.1. All weights were initialized using Xavier initialization(46). Model inputs were scaled by −100 and not normalized. The trained classifier and code for entropy calculations is available at the GitHub repository, *egurapha/prot_domain_segmentor.* The prerequisites are Python and the packages: PyTorch, Numpy, and SciPy. For nice visualizations, the PyMOL command line API is also recommended. All other data are available upon request.

### Rosetta

All *Rosetta* commands and *RosettaRemodel* blueprints are provided in *Supplemental Data*, available at https://goo.gl/osxG47.

## Acknowledgements

All data used in this study was obtained from the CATH protein database. A list of unique protein chains was kindly provided by Ian Sillitoe who works with the CATH group at University College London. Namrata Anand provided much technical advice, coding help, and shared many of her own tips for handling contact map data. This project was supported by startup funds from the Stanford Schools of Engineering and Medicine, the Stanford ChEM-H Chemistry/Biology Interface Predoctoral Training Program and the National Institute of General Medical Sciences of the National Institutes of Health under Award Number T32GM120007. Additionally, this project was supported by the U.S. Department of Energy, Office of Science, Office of Advanced Scientific Computing Research, Scientific Discovery through Advanced Computing (SciDAC) program.

## Author Contributions

R.R.E and P.-S.H. conceived and performed the research, and wrote the manuscript.

## Availability

The trained segmentor model and code for entropy calculation is available at the GitHub repository, *egurapha/prot_domain_segmentor.* The prerequisites are Python and the packages: PyTorch, Numpy, and SciPy. For visualization, the PyMOL command line API is also recommended. All other data are available upon request.

## Supplemental Information

### Supplemental Information

#### Model Extension for Multi-Domain Protein Classification

Conventional multi-domain protein classification entails two tasks. The first is *parsing*, in which sets of residues are grouped into domains. The second is *classification*, in which parsed domains are assigned a class label. While the model presented in the main text performs classification, it does not indicate which sets of residues comprise distinct domains. Because architecture classifications for any set of residues can be obtained by averaging over our classifier-predicted probabilities, our model can easily interface with previously reported domain parsing algorithms(1–4). Nonetheless, for completeness, we offer an additional domain-parsing CNN to augment this missing functionality. Probabilities predicted by the classifier network can be averaged over collections of residues specified by this new parser network to generate complete classifications **(Supplemental Figure S2)**.

Domain parsing can be formulated as a semantic segmentation problem where each residue is assigned to one of *k* classes each corresponding to a domain instance. Here, *k* is some maximum number of domains allowed by the model. (We choose *k*=*8*, although our dataset does not contain proteins comprised of more than five domains.) This task is similar to the classification approach described in the main text, with the only difference being that the model is required to identify the correct *sets* of residues as opposed to class labels. We therefore trained a model with an architecture identical to that of our classifier network, but with the output layer having 8 channels instead of 38 (**Figure 2a, Orange**), using the Hungarian algorithm(5) to compute set-difference losses during training.

The parser network converges within 30 epochs of training, achieving test accuracies of 91.6% per-structure, and 88.4% per-structure averaged over proteins with the same number of domains **(Supplemental Figure S3a)**. We achieve average accuracies higher than 80% even for proteins with more than four domains **(Supplemental Figure S3b)**. A confusion matrix for the predicted number of domains is shown in **(Supplemental Figure S3c)**. Example outputs are shown in **Supplemental Figure S4**.

To compare how our network performs against previous algorithms, we benchmarked our network on the Islam90(2, 6) and Jones(7) datasets that have been previously used to benchmark domain parsing algorithms. Our CNN is able to reproduce CATH domain parsing with accuracy comparable to that of previously reported methods (**Supplemental Table S5**).

---

**Figure S1:**
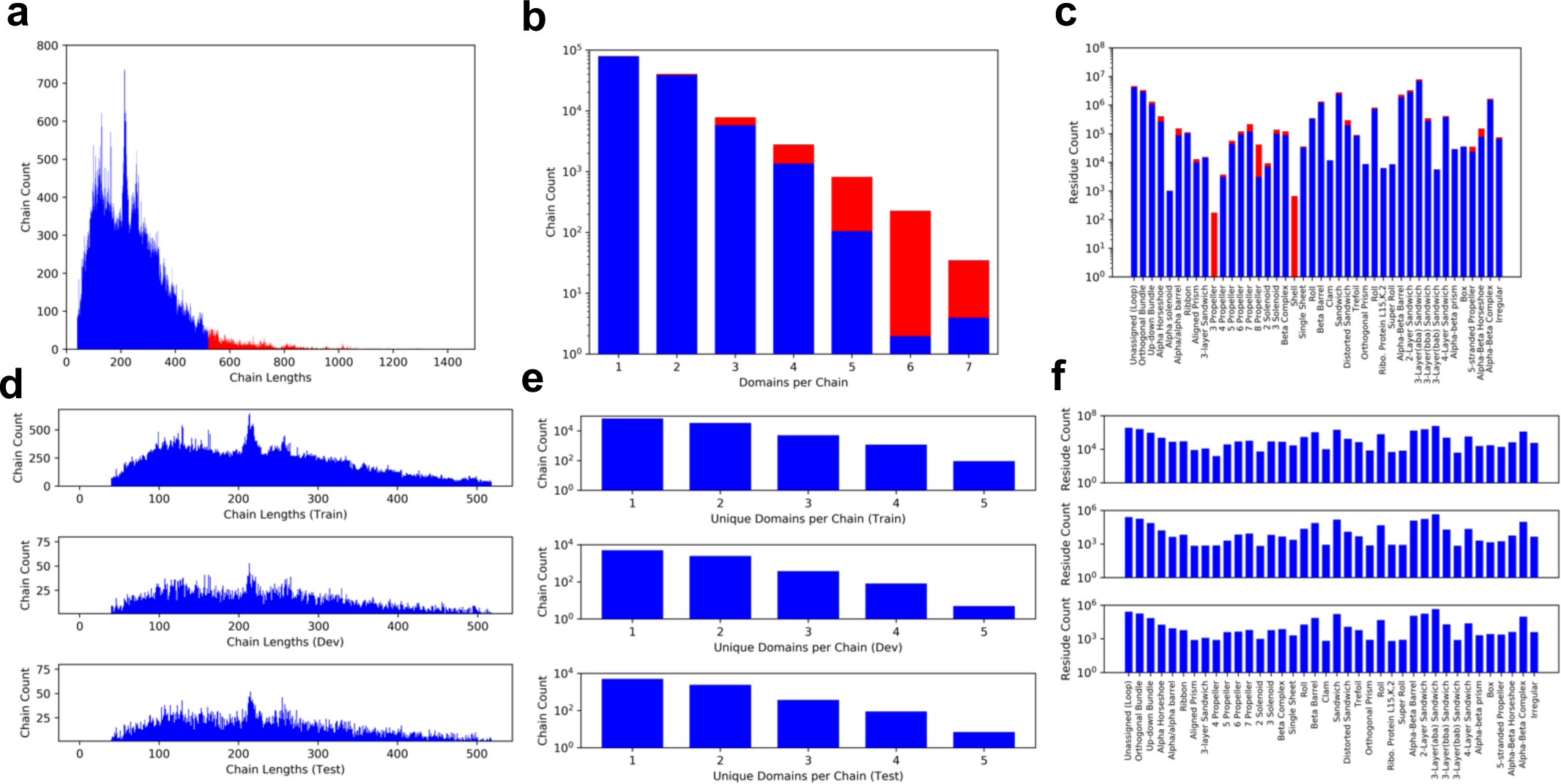
Dataset Structure. The distribution of data before and after selection are shown in **a, b**, and **c** which depict the distributions of chain lengths, number of domains per chain, and number of residues per class respectively. Selected data are shown in blue and unused data are shown in red. Overall the selection does not greatly alter the structure of the data obtained from CATH. The distribution of the selected data after splitting into training, development, and test sets is shown in **d**, **e**, and **f** which depict the distributions of chain lengths, number of domains per chain, and number of residues per class respectively. The split was performed in a stratified manner and stochastically adjusted to ensure that each class is represented by at least 650 residues in each set.

**Figure S2:**
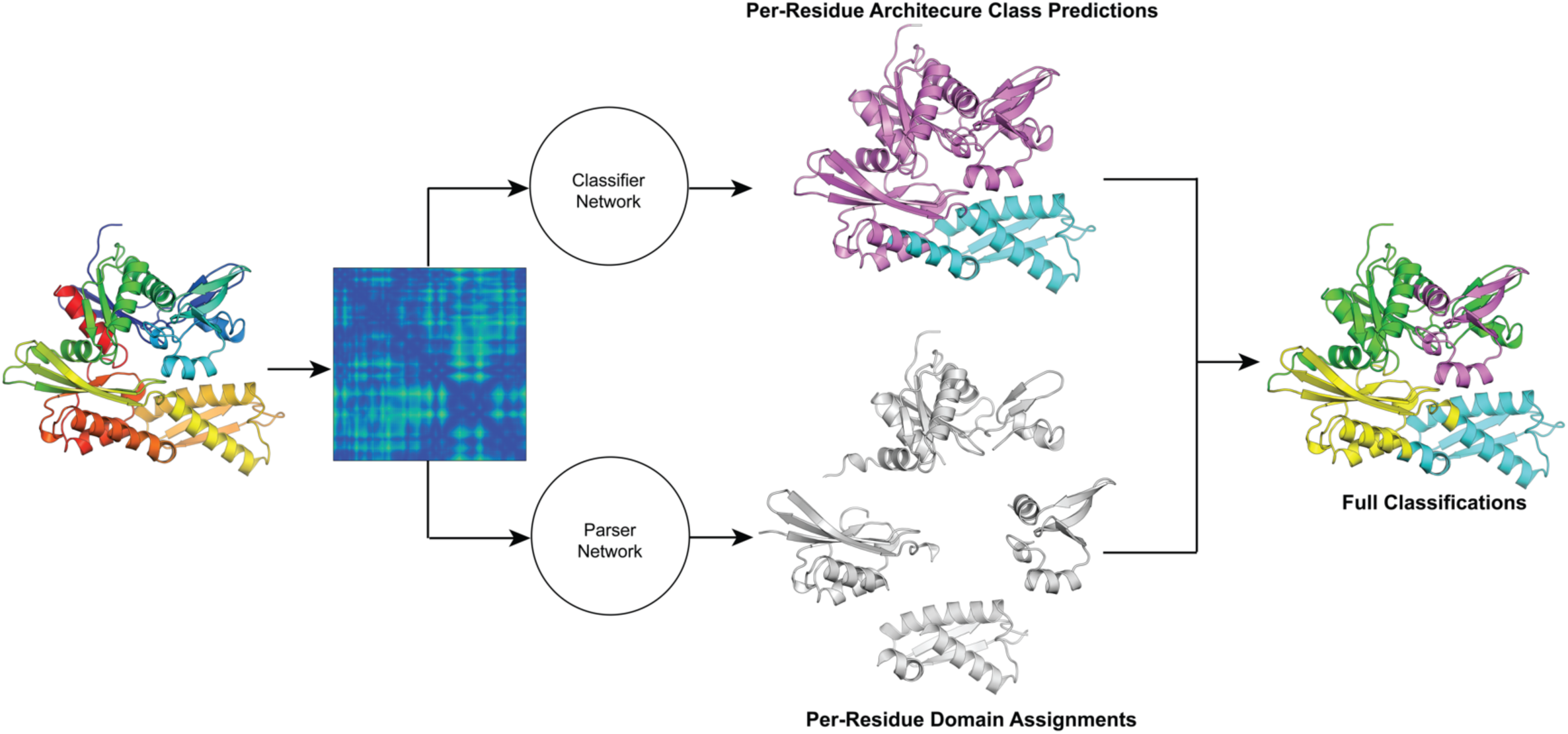
Schematic of a CNN Based Multi-Domain Protein Classifier. A schematic depicting how the classifier and domain parser networks can be combined to perform full classification of multi-domain proteins. Class assignments for residue sets specified by the parser network can be assigned an architecture by the classifier network (PDB ID: 5aqgE). The parser network can be replaced with previously reported domain parsing algorithms.

**Figure S3:**
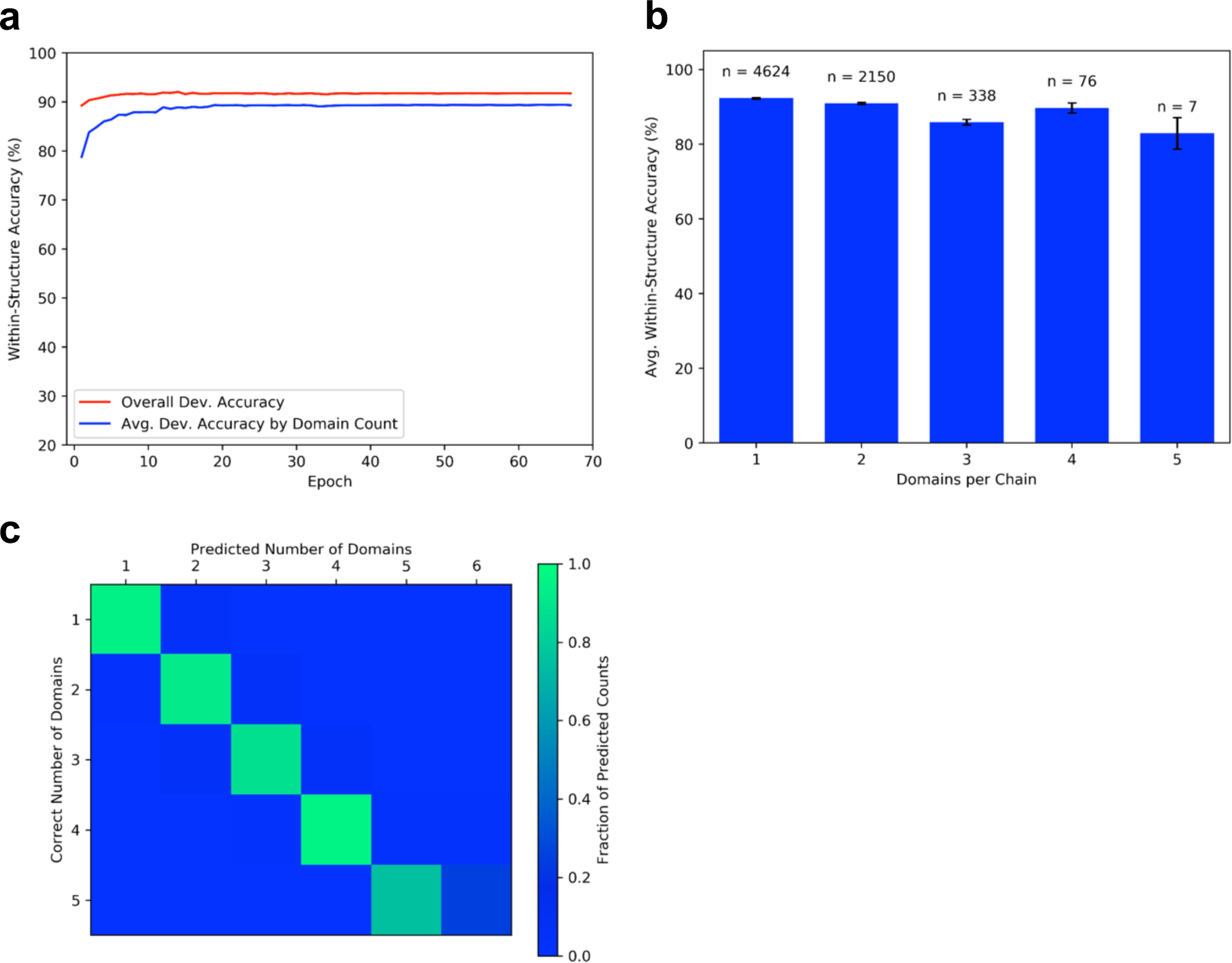
Parser Training and Performance. **a,** Training accuracy for the parser network. The red line indicates the average development accuracy per structure. The blue line indicates the per-structure development accuracy averaged across all proteins of the same length. The model converges within 30 epochs of training, achieving test accuracies of 91.6% per-structure, and 88.4% per-structure per-structure averaged over proteins with the same domain count. **b,** a bar graph showing the average per-structure test accuracies among structures with the same number of domains. Accuracies exceed 80% in all cases. The number of proteins with a given domain-count (*n*) is shown above each bar. **c,** a confusion matrix comparing the number of domains predicted by our model, and the number of actual domains in each test case.

**Figure S4:**
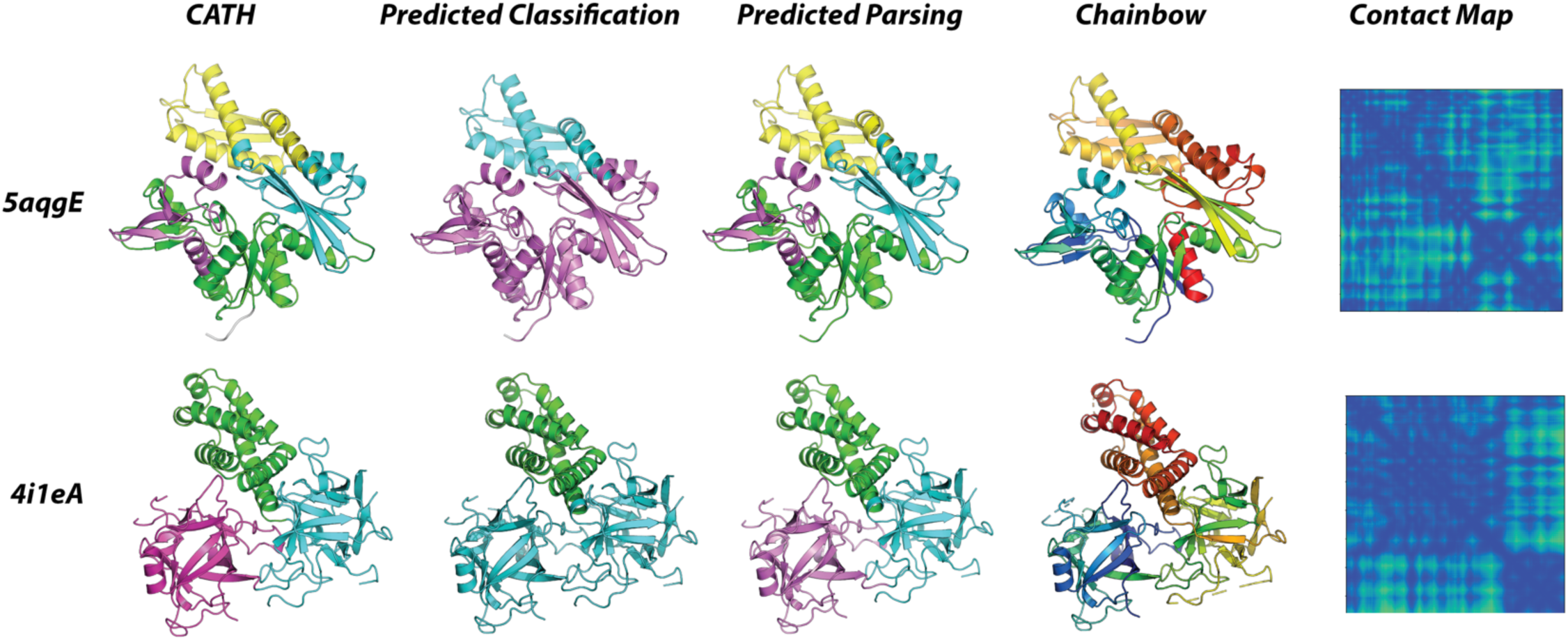
Multi-Domain Classification Examples. Example outputs from the classifier and parser network. Distinct domains in the predicted parsings are shown as unique colors. For both structures, our parser is able to closely reproduce CATH domain assignments. The 5aqgE structure is comprised of 4 domains, three of which belong to the αβ-complex class (5aqgE, predicted, magenta) and one of which belongs to the 2-layer sandwich class (5aqgE, predicted, cyan). The 4i1eA structure is comprised of 3 domains, two of which belong to the trefoil (4i1eA, predicted, cyan) and one of which belongs to the alpha-horseshoe class (4i1eA, predicted, green). The parser prediction in combination with per-residue classifications allow for full classifications.

**Table S5:**
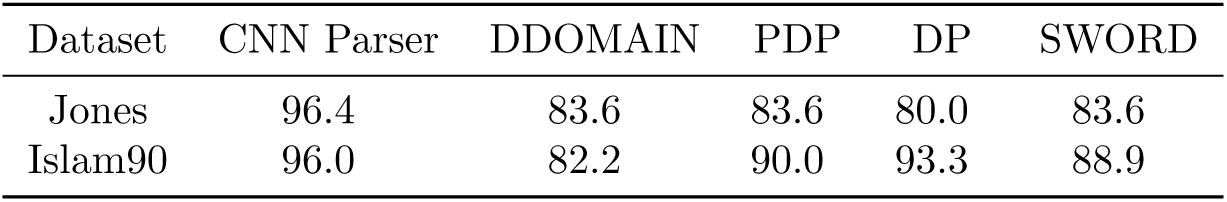
Parser Performance Benchmarks. A table comparing per-residue accuracies of our method (CNN Parser), Protein Domain Parser (PDP), Domain Parser (DP), DDOMAIN, and SWORD. Accuracy values are reported in percentages, and were obtained from the study by Postic et. al. (2017).

| Dataset | CNN Parser | DDOMAIN | PDP | DP | SWORD |
| --- | --- | --- | --- | --- | --- |
| Jones | 96.4 | 83.6 | 83.6 | 80.0 | 83.6 |
| Islam90 | 96.0 | 82.2 | 90.0 | 93.3 | 88.9 |

**Figure S6:**
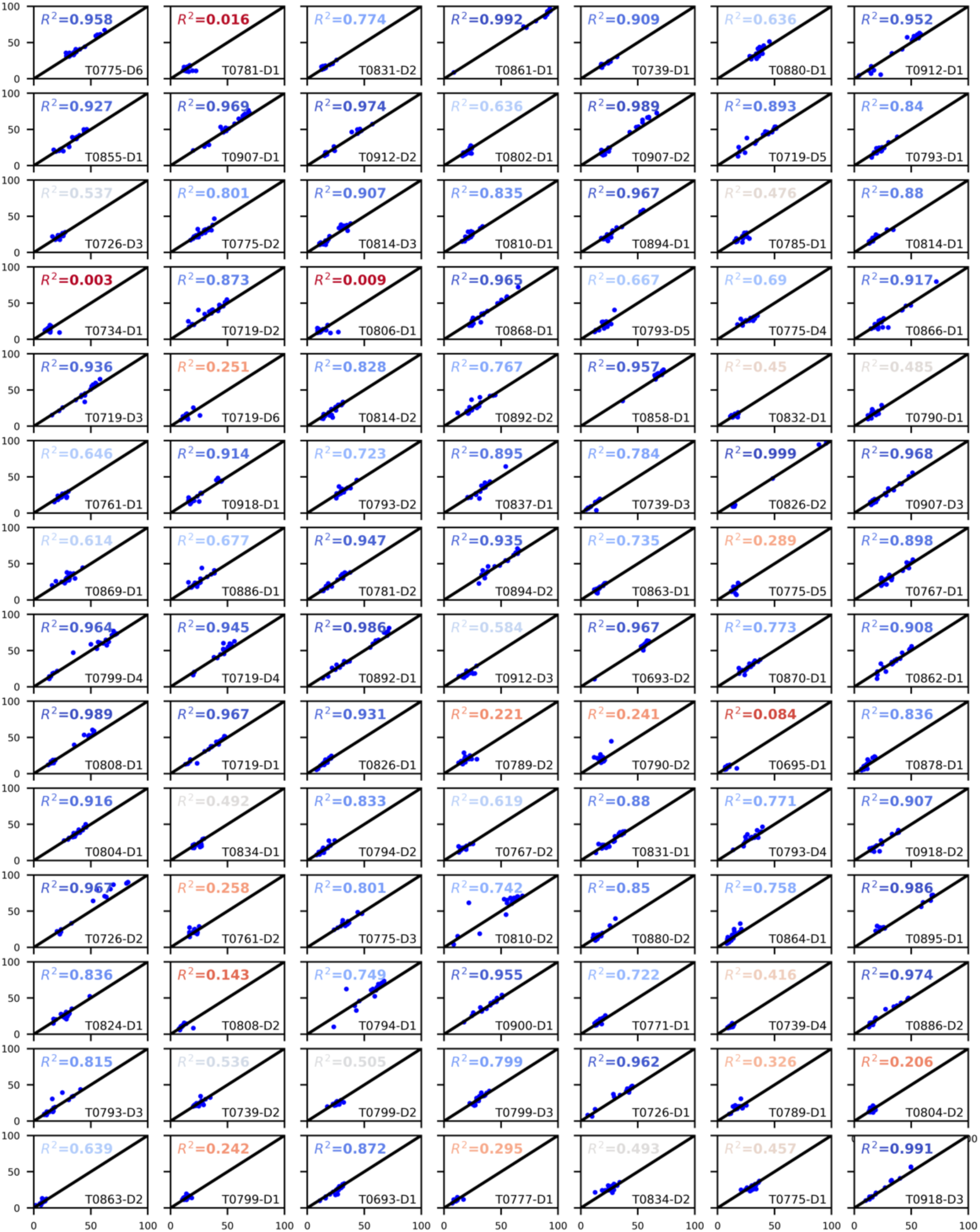
GDT-TS Score Prediction Profiles by CATH Target. Plots of real GDT-TS Values versus predicted GDT-TS values for each CATH target. The identity function is shown as a black line. Target names are shown in the bottom right corner of each plot, and R^2^ values are shown in the upper left.

